# Cortical oscillations reflect opponent ensemble dynamics through coordinated multifrequency activity

**DOI:** 10.64898/2026.02.20.707132

**Authors:** Jonathan Mishler, Morteza Salimi, Miranda Koloski, Irene Rembado, Carrie Shilyansky, Jyoti Mishra, Dhakshin Ramanathan

## Abstract

Neural oscillations are widely used as proxies for neuronal activity, where power in individual frequency bands is commonly interpreted as functionally indexing neural circuit engagement. However, power in individual frequency bands shows heterogeneous and sometimes opposing relationships with neuronal activity across regions and behavioral contexts, challenging the assumption of a stable frequency-to-circuit mapping. Here we show that glutamatergic population activity in rat medial prefrontal cortex is not stably linked with power in isolated frequency bands, but rather with dynamically recurring multi-frequency amplitude co-fluctuations. These multi-frequency patterns, termed spectral motifs, occurred in opponent pairs with nearly identical frequency composition but inverted relationships to population calcium activity. This opponent motif structure, observed across cortical regions and species, provides a key component for understanding how oscillations are linked to neuronal activity. We found that shifts in motif opponency balance explained changes in glutamatergic activity that occur during brain-computer interface learning better than models based on frequency band power alone. Furthermore, opponent motifs map selectively onto opponent cell ensembles and enable bidirectional mapping between local field potentials and ensemble activity. These findings identify multi-frequency opponent motifs as a conserved organizational principle linking oscillatory dynamics to population-level circuit states and challenge the notion that individual frequency bands can serve as interpretable functional units mapping onto neural circuit activity.

## Main

Neuronal spike activity provides the most direct and temporally precise measure of circuit output^1,2^, capturing both the activity of individual neurons and their coordination at the population level^3,4^. In contrast, oscillatory signals such as local field potentials (LFPs) reflect aggregate synaptic and dendritic currents from local neuronal populations^5,6^, offering a complementary view of circuit dynamics. Neural oscillations are among the most prominent and widely observed features of brain activity across species and recording modalities^6,7^ and have therefore been widely treated as a bridge between microscale circuit activity and large-scale network organization^8,9^. However, the relationship between oscillatory activity and neural or behavioral variables is often heterogeneous and context dependent^10–12^. Complicating their relationships, power within a given frequency band has also been shown to correlate positively or negatively with spiking within the neocortex^13^. These heterogeneous relationships challenge the assumption that individual frequency bands provide stable and interpretable mappings onto specific circuit states. While certain structured cross-frequency phenomena, including nested rhythms and phase–amplitude coupling, have been linked to identifiable circuit interactions in specific brain circuits^14,15^, it remains unclear whether multi-frequency organization more generally structures the relationship between oscillatory dynamics and single cell/population activity. As a result, there is still no consistent framework for relating specific neuronal ensembles to specific patterns of oscillatory activity across contexts.

A central challenge in linking oscillations to spike activity arises from their distinct biophysical origins. LFPs primarily reflect summed transmembrane currents dominated by synaptic and dendritic processes^5,16^, whereas spikes report only suprathreshold neuronal output^1,2^. We hypothesized that calcium imaging may provide a critical bridge between these signals by capturing both suprathreshold firing and subthreshold integrative dynamics across genetically defined neuronal populations^17^, enabling direct comparison between ensemble activity and multiband oscillatory structure. Here, we combined LFP recordings with either bulk fiber photometry or single-cell calcium imaging in rat medial prefrontal cortex (mPFC) to measure oscillatory dynamics alongside glutamatergic activity. We quantified amplitude co-fluctuations across frequency bands and their relationship to bulk calcium signals to identify opponent spectral motifs. We then examined the behavioral relevance of these motifs in a closed-loop brain-computer interface (BCI) task and their relationships to single-neuron and ensemble calcium activity. Finally, we tested whether similar opponent multi-frequency structures were detectable across cortical regions and in noninvasive human electroencephalography (EEG) recordings.

### Multi-frequency spectral motifs reveal opponent structure in cortical oscillations

Within mPFC, smoothed LFP amplitudes across frequency bands (1-80 Hz) appeared to exhibit structured, frequency-specific relationships with glutamatergic population calcium activity (Figure 1C). Relationships were quantified as the Pearson correlation coefficient computed across time between each band-limited LFP amplitude envelope and the calcium trace. We repeated the analysis after circularly shifting the calcium trace by 1200 s to remove time alignment between the signals, which attenuated these calcium-frequency correlations. Correlations were consistent across animals and sessions (N = 6 rats, 3 sessions per rat) and were robust to smoothing time and bandwidth selection (Supplementary Figure 1). However, on closer inspection of temporal dynamics, we observed that these frequency-calcium relationships were neither stationary nor independent. Sliding-window analyses revealed that correlations between individual frequency bands and calcium activity varied over time and frequently reversed sign (Figure 1D,E). Moreover, correlations across frequency bands did not fluctuate independently but instead exhibited coordinated higher-order structure (Figure 1E). Thus the average correlations shown in Figure 1C relied on assumptions of stationarity and independence that do not reflect these neural dynamics.

**Figure 1.**
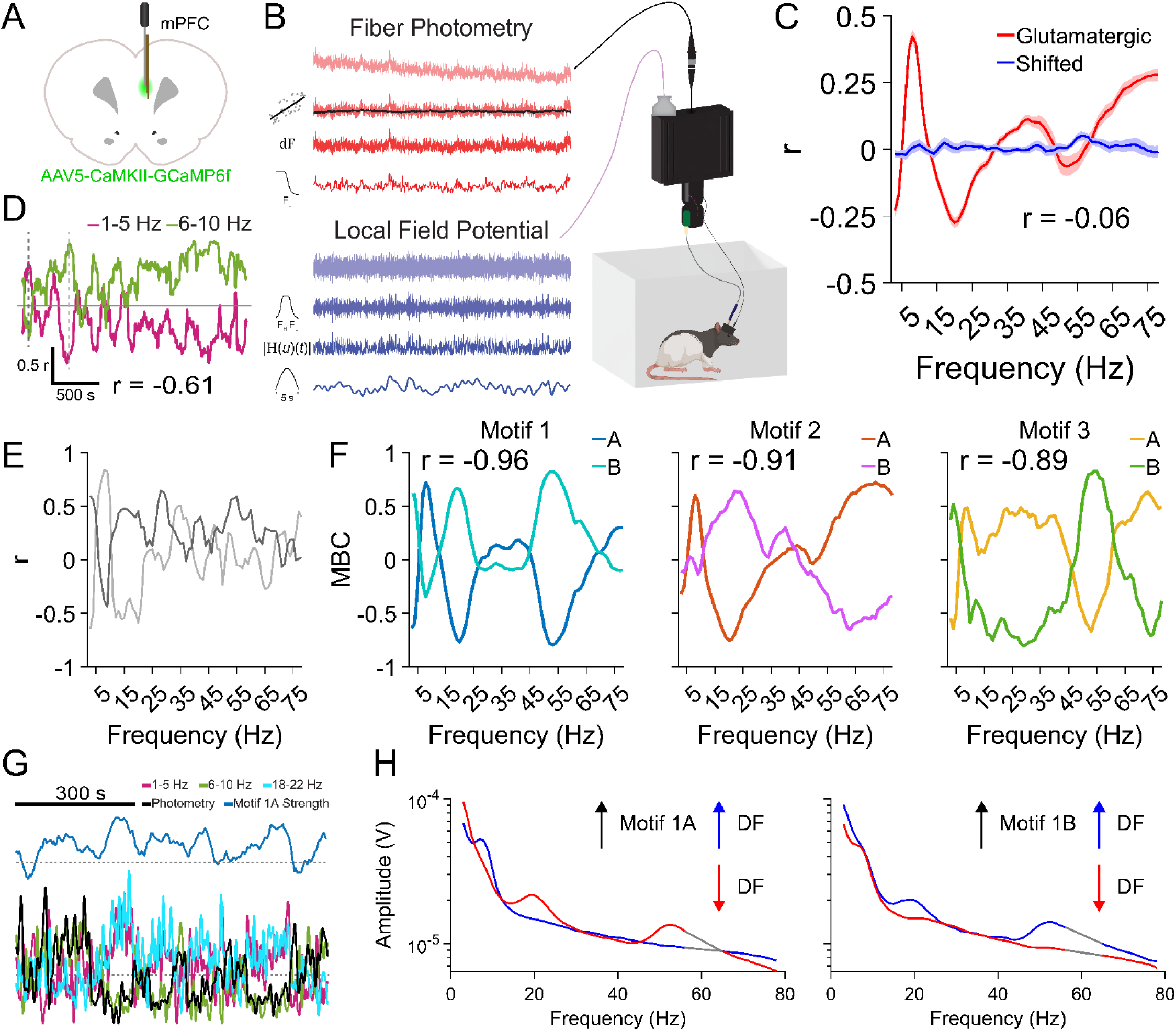
Opponent spectral motifs structure the relationship between LFP amplitude and glutamatergic population activity. (A) Custom-fabricated optrode implantation in rat mPFC. (B) Recording setup and preprocessing of calcium and LFP signals. (C) Frequency-specific correlations between LFP amplitude and calcium activity. Pearson correlation coefficients were computed across time between each band-limited amplitude envelope and the calcium trace (red). Circularly shifting the calcium trace by 1200 s to disrupt temporal alignment attenuated both correlation magnitude and spectral structure (blue). Shaded regions denote ± s.e.m. across sessions (N = 6 rats, 3 sessions per rat). (D) Sliding-window correlations reveal non-stationary relationships between individual frequency bands and calcium activity. (E) Cross-frequency correlations of sliding-window correlation coefficients demonstrate coordinated multiband structure rather than independent fluctuations across frequencies. (F) Identification of recurring multiband opponent spectral motifs. (G) Representative 800 s segment illustrating coordinated amplitude co-fluctuations across selected frequency bands (1–5 Hz, 6–10 Hz, 18–22 Hz) during expression of Motif 1 and its relationship to calcium activity. (H) Average LFP amplitude across frequencies during periods of high Motif 1A versus Motif 1B expression. Identical frequency bands index opposing calcium states depending on opponent motif dominance. Grey lines indicate band-stopped frequencies.

To characterize this cross-frequency organization with greater temporal resolution than afforded by traditional correlational analyses (which rely on assumptions of stationarity), we transformed the band-limited amplitude envelopes into a time-resolved measure of frequency-specific directional concordance with population calcium activity (see Methods). At each time point, z-scored smoothed amplitudes from all frequency bands were multiplied by the z-scored calcium signal, and the sign of this product was taken to yield a binary concordance measure (BCM) for each frequency band, reflecting whether LFP amplitude and calcium activity were aligned or opposed at that moment. The resulting multi-dimensional BCM vector defined a multi-frequency directional state at each time sample. Time points were grouped according to similarity in these cross-frequency concordance patterns, allowing identification of recurring spectral motifs expressed throughout the recording. To ensure robustness and generalizability, we required that motifs be reproducible across animals and sessions. This procedure yielded a small set of spectral motifs in mPFC (Figure 1F), each defined by a characteristic multi-frequency pattern of directional concordance with population calcium activity. Notably, each motif occurred as an opponent pair (labeled A and B in figure), with nearly identical frequency composition but opposite relationships to population calcium activity. Opponent motifs occurred at distinct times during the recordings and were consistent with alternating network states in which the same multi-frequency amplitude configuration was associated with either elevated or suppressed glutamatergic population activity. Both motif structure and opponency were independently recovered using PCA and factor analysis (Supplementary Figure 2), suggesting they were not artifacts of our analytic method. Importantly, these spectral motifs were also directly observable in the filtered data rather than arising solely from latent projections. A representative 800 s segment illustrates coordinated amplitude co-fluctuations across 1-5 Hz, 6-10 Hz, and 18-22 Hz bands associated with Motif 1 and their relationship to calcium activity (Figure 1G).

The presence of opponent spectral motifs indicates that the relationship between amplitude within any individual frequency band and population activity is dynamic. To illustrate this, we compared average LFP amplitudes during periods of high Motif 1A versus Motif 1B expression (Figure 1H). During high Motif 1A expression, elevated calcium activity coincided with increased theta-band amplitude and reduced delta, beta, and gamma amplitudes, whereas the inverse pattern occurred during Motif 1B expression. Thus, identical frequency bands indexed opposite population calcium states depending on opponent motif dominance, demonstrating that single band-limited LFP amplitude alone does not provide a stable or monotonic readout of glutamatergic population activity, which is instead interpretable from the perspective of the multi-frequency opponent motif structure.

### Volitional modulation of opponent spectral motif expression

To test whether opponent spectral motifs were behaviorally relevant rather than epiphenomenal, we trained rats on an auditory, trial-based BCI task in which population calcium activity was coupled to an auditory cursor (Figure 2A). In resting-state recordings, elevated expression of Motif 1A was associated with increased calcium activity, whereas elevated expression of the opposing Motif 1B was associated with decreased calcium activity (Wilcoxon signed-rank test, Motif 1A p=0.007, median=0.077; Motif 1B p=2 x 10^-4^, median=-0.25; Figure 2B). No other motif showed a consistent relationship with increases or decreases in calcium activity. Based on this dissociation, we hypothesized that animals trained to suppress calcium activity would preferentially recruit/activate Motif 1B and lower recruitment of the opponent Motif 1A during the BCI trials. The BCI task involved real-time processing of bulk calcium activity, averaged in 100 ms bins and compared to a fixed threshold set for that animal/session that dynamically adjusted across trials. Bins below threshold increased the pitch of an auditory cursor toward the rewarded target frequency, whereas bins above threshold decreased it (Figure 2C; see Methods for details).

**Figure 2.**
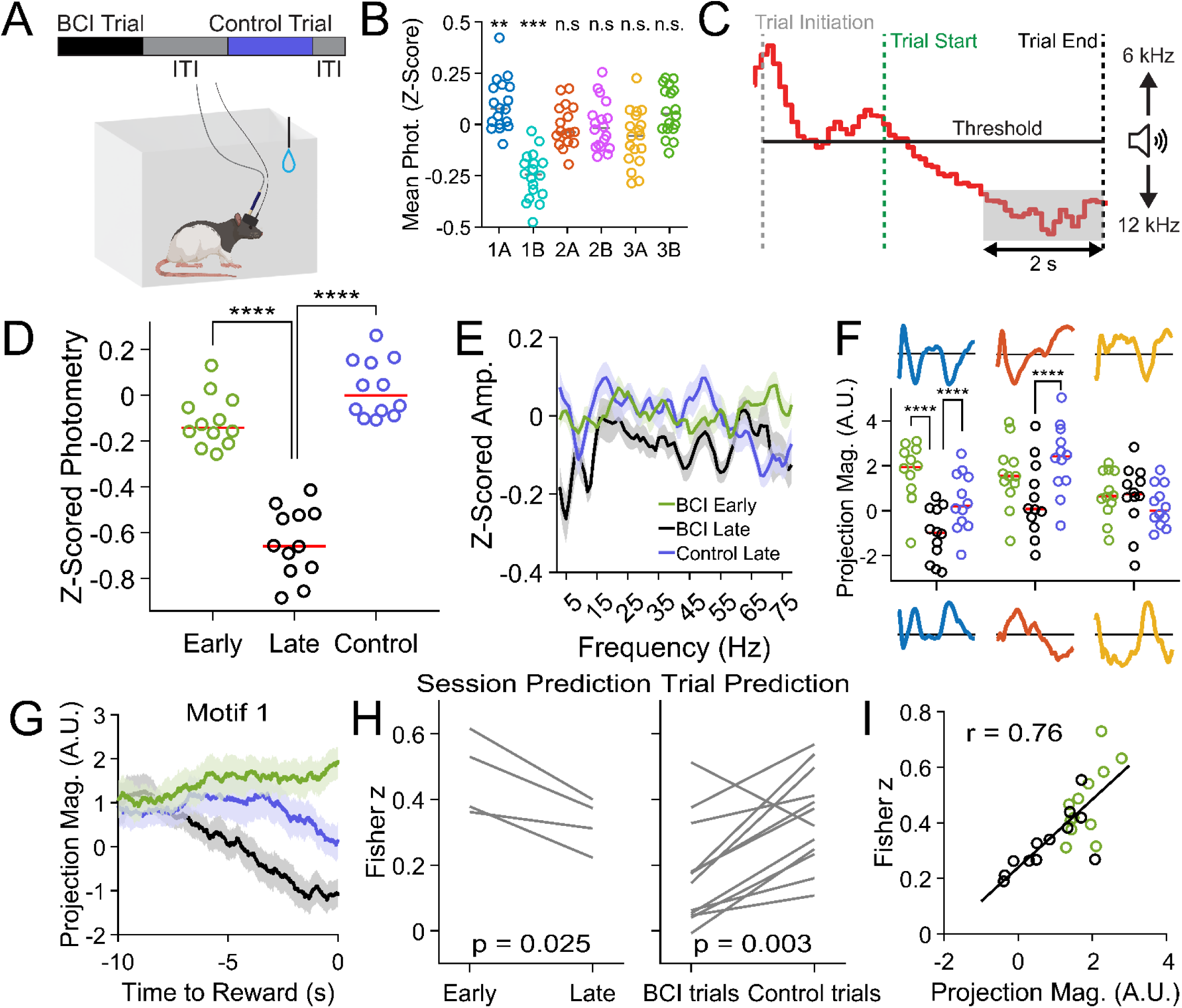
Volitional modulation of opponent spectral motif expression during brain-computer interface learning. (A) Schematic of the auditory BCI task linking population calcium activity to an auditory cursor. (B) In resting-state recordings, elevated Motif 1A expression was associated with increased calcium activity, whereas Motif 1B expression was associated with decreased calcium activity (Wilcoxon signed-rank tests, p < 0.01). (C) Trial structure and thresholding logic for closed-loop control of the auditory cursor, in which 100 ms bins of calcium activity were compared to a fixed threshold to modulate pitch. (D) Population calcium activity decreased during BCI learning. Calcium levels were significantly reduced during late learning relative to early learning and control trials (Welch and paired t-tests, both p < 10⁻⁸), confirming robust learning-dependent suppression of population activity. (E) Late learning was accompanied by decreases in LFP power across multiple frequency bands. (F) Learning-dependent increases in Motif 1B expression were observed during late BCI trials relative to early learning and control conditions (Welch t-test and paired t-test, both p < 10⁻⁴). (G) Trial-aligned analyses revealed selective elevation of Motif 1B expression during task engagement in late learning. (H) Linear model trained on resting-state LFP generalized poorly during late learning and BCI trials, when Motif 1B expression was elevated. (I) Across sessions, decoding performance scaled with the relative balance of time-averaged projection strength of A and B opponent motifs (Pearson r = 0.76, p = 1.4 × 10⁻⁵).

Following training, rats reliably reduced population calcium activity during task performance relative to both early learning and control trials (early vs late learning: Welch *t*(20.37) = 9.70, *p* = 4.40 × 10⁻⁹, Hedges *g* = −3.83; late learning vs control: paired *t*(11) = −42.75, *p* = 1.40 × 10⁻¹³, dz = −12.34; nonparametric checks: ranksum *p* = 3.66 × 10⁻⁵, sign-rank *p* = 4.88 × 10⁻⁴; Figure 2D). These reductions were accompanied by decreases in LFP power across multiple frequency bands (Figure 2E), consistent with a global shift in oscillatory structure rather than modulation of a specific band. Critically, as hypothesized, calcium modulation was associated with a selective increase in Motif 1B expression during late BCI trials compared to both early learning and control conditions (early vs late BCI trials: unpaired Welch *t*(21.7) = 5.35, *p* = 2.33 × 10⁻⁵, FDR-corrected *p* = 7.00 × 10⁻⁵, Hedges *g* = −2.11; late BCI trials vs control trials: paired *t*(11) = −15.84, *p* = 6.42 × 10⁻⁹, FDR-corrected *p* = 1.35 × 10⁻⁸, dz = −4.57; Figure 2F). Other motifs did not show learning-dependent changes. Trial-aligned analyses further confirmed that Motif 1B expression increased specifically during task engagement in late learning, rather than reflecting nonspecific session-level changes (Figure 2G).

These results indicate that expression of opponent spectral motifs best explains the changes in the oscillatory dynamics that occur during the BCI task. To probe this further, we trained linear models to predict calcium activity from resting-state LFP (using a monotonic mapping of frequencies to calcium and ignoring motif structure), and analyzed their performance on data collected during early and late sessions of BCI task performance. If motif structure was unimportant, BCI task performance should not disrupt overall model performance, whereas if motif opponency structure was important, the model performance should diminish. Consistent with this framework, linear models trained to predict calcium activity from resting-state LFP generalized poorly during late learning sessions and BCI trials when Motif 1B expression was elevated (Figure 2H). Across sessions, prediction performance scaled with the relative balance of time-averaged concordance projection strength of A and B opponent motifs (Pearson r = 0.76, p = 1.4 × 10⁻⁵; Figure 2I), indicating that decoding failures arose from shifts in opponent motif dominance rather than changes in noise or signal quality.

### Spectral motifs map onto opponent neuronal ensembles

We identified two important properties of LFP oscillations: multi-frequency patterns of activity (motifs) and motif opponency (opposite relationships with bulk calcium activity). We hypothesized that these would be represented within single cell activity. If true, this would entail two predictions: 1) single cells would be correlated with multi-frequency motifs, and 2) opponent motifs would be reflected by distinct populations of cells. A second hypothesis is that motifs are not reflected by single cell activity, but rather that single cells are linked with individual frequencies and motifs reflect the summed population or ensemble activity (e.g. distinct populations of cells that, together, reflect the complex frequency patterns). To test this, we simultaneously recorded LFP and single-neuron calcium activity using a one-photon miniaturized microscope (miniscope). We first asked whether average calcium-LFP and motif-level relationships identified using fiber photometry generalized to single-cell recordings. Mean population calcium activity extracted from one-photon imaging exhibited frequency-specific correlations with LFP amplitudes that closely matched those observed with fiber photometry (r = 0.83; Figure 3A), indicating that the correlational structure was preserved across recording modalities and processing methods (see Methods for detailed differences).

**Figure 3.**
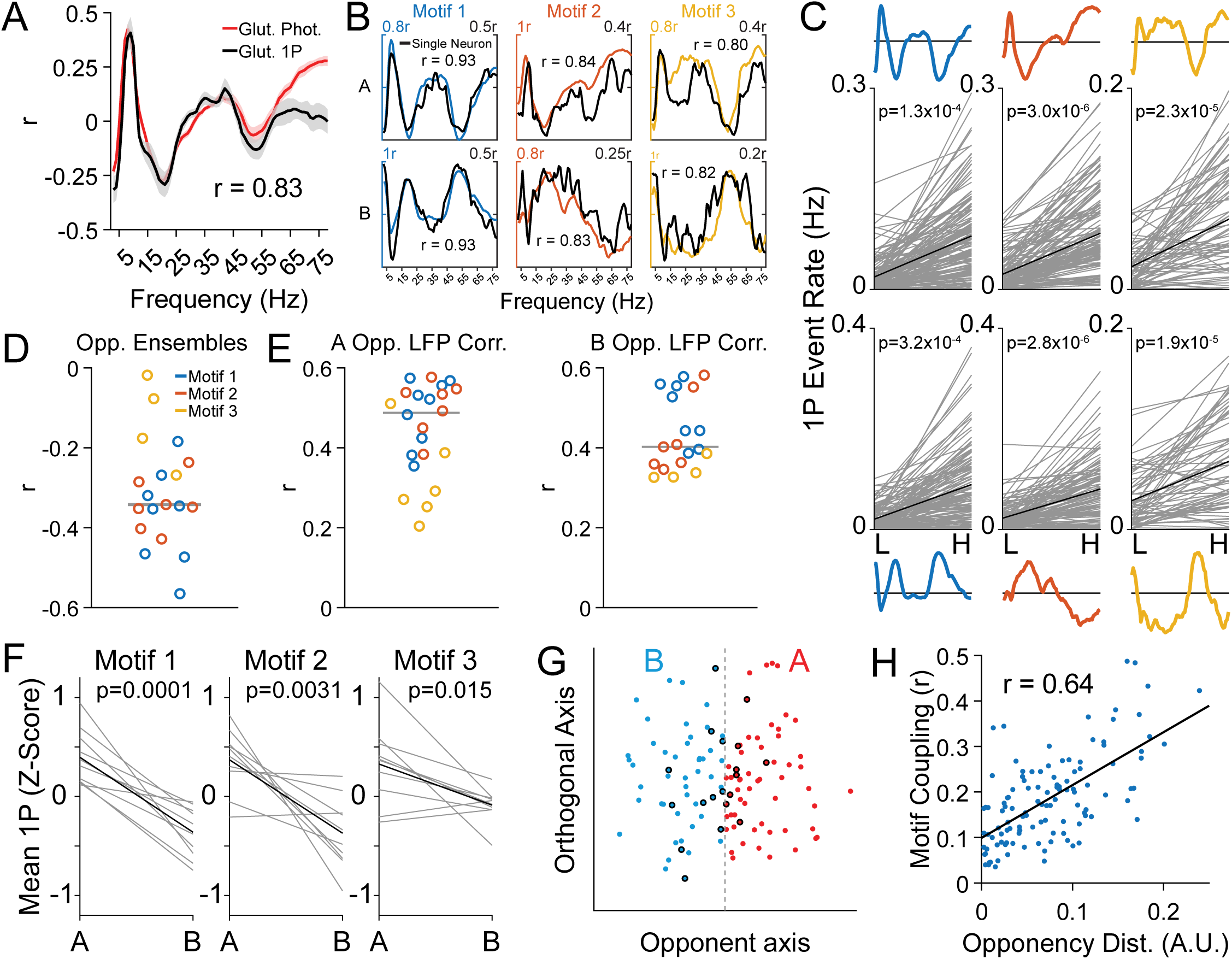
Opponent spectral motifs map onto opponent neuronal ensembles. (A) Frequency-specific correlations between mean one-photon population calcium activity and LFP amplitudes closely matched those observed with fiber photometry (r = 0.83). (B) Individual neurons exhibited multiband LFP correlation profiles aligned with opponent spectral motif templates. (C) Neurons assigned to a given opponent motif increased spiking when LFP activity projected onto that motif and decreased spiking when the opposing motif dominated. Spike rates covaried significantly with projection strength of the corresponding motif (paired t-tests across sessions, all p < 10⁻³). (D) Ensembles defined by opposing components of the same motif exhibited significant anticorrelations in mean spike rates across sessions for Motifs 1 and 2 (Motif 1: p = 6.0 × 10⁻⁵; Motif 2: p = 8.7 × 10⁻⁶; one-sample t-tests; Motif 3: p = 0.09). (E) Ensemble spike rates covaried with projection strength of the corresponding LFP motif template across all opponent motifs (one-sample t-tests across sessions, all p < 10⁻⁴).(F) Bulk calcium activity reflected opponent ensemble balance. Periods dominated by A motifs were associated with higher mean calcium activity than periods dominated by corresponding B motifs (paired t-tests across sessions; Motif 1 p = 0.0001; Motif 2 p = 0.0031; Motif 3 p = 0.015). (G) Principal component analysis of z-scored single-cell calcium traces revealed an opponent axis separating neurons into two opponent groups. (H) Neurons more strongly aligned with a given opponent motif exhibited greater separation along the opponent axis (paired t-test, p = 0.037).

We next examined relationships between individual neurons and LFP frequency bands. To test this prediction, we quantified moment-to-moment motif expression using LFP projections (see Methods) by binarizing the amplitude of each frequency band by calculating the sign of its z-scored amplitude (positive vs negative deviation) and projecting these bandwise binary vectors onto each opponent motif template. Correlating each neuron’s calcium trace with LFP amplitudes across frequency bands revealed that cells exhibited multi-frequency correlation profiles that closely matched those defined by the opponent spectral motifs (Figure 3B). Because each opponent motif is characterized by a specific multi-frequency amplitude pattern relative to population calcium activity, a neuron whose frequency-specific correlation profile aligns with a given motif should increase its activity when the LFP expresses that motif’s pattern and decrease its activity when the opposing pattern dominates. We assigned each neuron to the opponent motif whose LFP projection time series showed the strongest correlation with its calcium activity. Consistent with this motif-defined structure, neurons assigned to a given opponent motif exhibited increased calcium spiking when LFP activity projected strongly onto that motif template and reduced spiking when projection favored the opposing template. Spike rates covaried significantly with projection strength of the corresponding motif (paired t-tests across sessions, all p < 10⁻³; Figure 3C), allowing motif expression in the LFP alone to predict single-cell ensemble activity. This suggests that motif opponency is itself reflected by opponency represented within the ensembles themselves.

To explore this, we averaged spike rates across neurons within each session and assessed the relationships between resulting ensemble activities. Ensembles corresponding to opposing components of the same motif exhibited significant anticorrelations in their mean spike rates for Motifs 1 and 2 (one-sample t-tests across sessions; Motif 1: p = 6.0 × 10⁻⁵; Motif 2: p = 8.7 × 10⁻⁶; Figure 3D). Although Motif 3 did not reach significance at the ensemble level (p = 0.09), Motif 3 opponent ensembles displayed consistent negative correlations across all sessions. These results indicate that the spectral motifs identified opponent neuronal ensembles with coordinated population dynamics. Next, to directly link ensemble activity to motif expression, we correlated ensemble spike rates with LFP projection strength for each opponent motif. Across all opponent motifs, ensemble spike rates covaried significantly with LFP projection strength (one-sample t-tests across sessions, all p < 10⁻⁴; Figure 3E), such that strong expression of a given opponent motif was accompanied by increased spiking in the corresponding ensemble and reduced spiking in its opponent. We then tested whether motif-level LFP structure captured ensemble activity more effectively than any individual frequency band in isolation. For each opponent ensemble, correlations between ensemble spike rates and motif projection strength exceeded those between ensemble spike rates and any single frequency band (76 frequency bands × 6 opponent motifs = 456 tests), with 89.5% reaching statistical significance after correction for multiple comparisons. Thus, across all analyses, we found that at the ensemble level, multi-frequency spectral motifs provide a more faithful representation of population spiking activity than any individual LFP frequency band.

### Opponent ensemble organization is functional rather than anatomical

Having established that opponent spectral motifs map onto opponent neuronal ensembles, we next examined how these ensemble dynamics are reflected in bulk calcium activity. Periods of strong LFP projection onto either opponent motif were associated with increased spiking in the corresponding ensemble, yet bulk calcium fluorescence exhibited a consistent asymmetry: periods dominated by A motifs were associated with higher mean calcium activity, whereas periods dominated by the corresponding B motifs were associated with significantly lower mean calcium activity across all motifs (paired t-tests across sessions; Motif 1 p = 0.0001; Motif 2 p = 0.0031; Motif 3 p = 0.015; Figure 3F). Thus, bulk calcium activity reflects the relative balance between opponent ensembles rather than providing a symmetric readout of ensemble activation.

If opponent ensemble structure reflects intrinsic population organization, it should be recoverable from calcium activity alone, independent of LFP measurements. Principal component analysis of z-scored single-cell calcium traces revealed a separable opponent axis along which neurons could be divided into two groups (Figure 3G). Assigning these groups to A or B components based on their relationship to bulk calcium activity enabled recovery of opponent ensembles corresponding to Motifs 1–3, with a median agreement of assignment rate of 72% across sessions. Neurons more strongly aligned with a given opponent motif also exhibited greater separation along the opponent axis in calcium space (paired t-test, p = 0.037; Figure 3H). Beyond opponency, we did not identify consistent physiological features distinguishing motifs, including firing rates or interspike interval statistics (Supplementary Figure 3), indicating that ensemble separation was driven by opponent structure rather than gross firing properties. Finally, to determine whether opponent motif-defined ensembles reflected anatomical segregation rather than functional organization (for example, cells that are closer together or within a similar cortical layer are more likely to be associated with a particular motif), we examined their spatial arrangement within the local cortical circuit. Nearest-neighbor analyses revealed minimal deviation from chance organization, with small effect sizes and few significant motif-pair comparisons relative to shuffled controls (97.6% of permutations p>0.05). These results indicate that opponent neuronal ensembles are spatially intermingled rather than anatomically clustered within mPFC (Supplementary Figure 4).

### Opponent spectral motifs enable bidirectional mapping between cellular and LFP activity

To determine whether opponent motif structure was sufficient to link cellular activity to multi-frequency LFP dynamics, we reconstructed smoothed LFP amplitude across frequency bands from single-cell calcium activity. Each neuron’s calcium trace was weighted by the frequency mapping associated with its assigned opponent motif, and the resulting signals were summed across neurons to generate reconstructed LFP power (Figure 4A). Reconstruction accuracy varied systematically across frequencies, with higher correlations at frequencies strongly represented in the spectral motifs and minimal reconstruction at weakly represented frequencies (Figure 4B, black). This frequency dependence suggested that reconstructable LFP structure was constrained by motif-level spectral organization (Figure 4B, grey).

**Figure 4.**
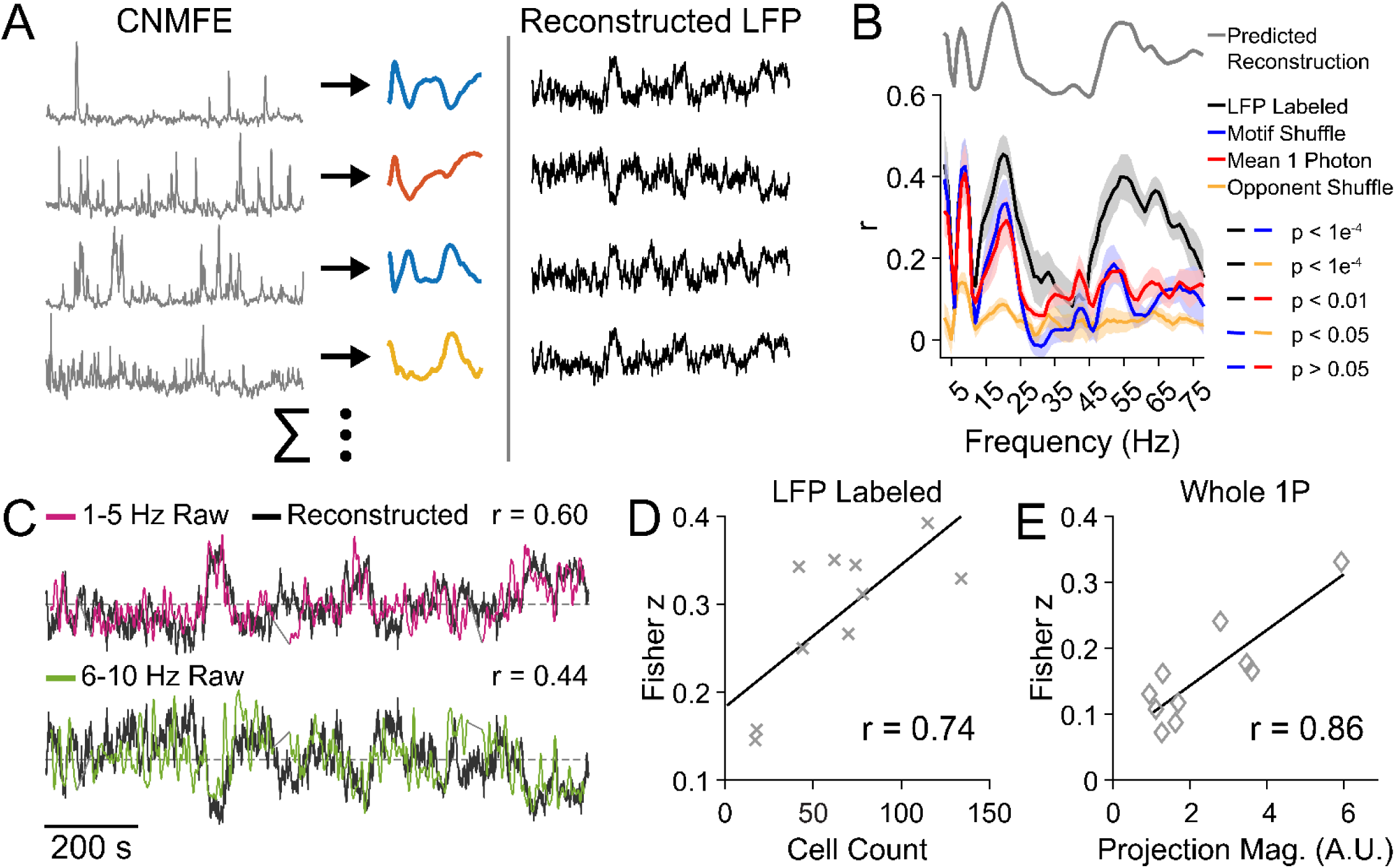
Motif identity and opponency are required for bidirectional reconstruction of multiband LFP dynamics from cellular activity. (A) LFP power reconstruction: each neuron’s calcium trace was weighted by the frequency mapping associated with its assigned opponent motif and summed across neurons to generate reconstructed band-limited LFP amplitude. (B) Reconstruction performance across conditions. Reconstructions preserving both motif identity and opponency (black) significantly outperformed motif-shuffled (blue), opponent-shuffled (orange), and mean 1-photon calcium (red) controls (paired t-tests on Fisher z–transformed correlations across sessions; labeled vs motif-shuffled: p = 4.42 × 10⁻⁵; labeled vs opponent-shuffled: p = 2.93 × 10⁻⁵; labeled vs mean 1P calcium: p = 0.0033). Motif-shuffled reconstructions also significantly outperformed opponent-shuffled reconstructions (p = 0.0116), indicating that disrupting opponency produced a larger degradation in reconstruction accuracy than disrupting motif identity. (C) Representative 20-minute traces of LFP amplitude in two frequency bands (1–5 Hz and 6–10 Hz, magenta and green, respectively) and corresponding reconstructions (black) without shuffling. (D) Reconstruction accuracy increased with the number of neurons recorded per session when motif and opponent identity were preserved (Pearson r = 0.74, p=0.015). (E) Reconstruction based on mean 1P calcium activity scaled with dominance of A motifs (Pearson r = 0.86, p = 0.0014).

To directly test the necessity of motif identity and opponency, we compared reconstruction performance against three control conditions: motif-shuffled reconstructions preserving opponency but randomizing motif identity (blue), opponent-shuffled reconstructions preserving motif identity but randomizing opponency (orange), and reconstructions based on mean 1-photon calcium activity alone (red). Reconstructions preserving both motif identity and opponency significantly outperformed all control conditions across sessions (paired t-tests on Fisher z–transformed correlations; labeled vs motif-shuffled: p = 4.42 × 10⁻⁵, FDR-corrected p = 8.84 × 10⁻⁵, dz = 2.32; labeled vs opponent-shuffled: p = 2.93 × 10⁻⁵, FDR-corrected p = 8.84 × 10⁻⁵, dz = 2.44; labeled vs mean 1P calcium: p = 0.0033, FDR-corrected p = 0.0044, dz = 1.25; Figure 4B). In addition, motif-shuffled reconstructions significantly outperformed opponent-shuffled reconstructions (paired t-test, p = 0.0116, FDR-corrected p = 0.0116, dz = 1.00), indicating that disrupting opponency produced a larger degradation in reconstruction accuracy than disrupting motif identity. Motif-shuffled reconstructions showed selective degradation in beta and gamma frequency ranges where motif-specific spectral structure diverged most, whereas opponent-shuffled reconstructions exhibited broadband degradation across frequencies. These findings reveal a hierarchical structure in which motif identity refines frequency-specific structure, but opponency provides the primary constraint shaping the relationship between LFP dynamics and population activity. Representative 20-minute traces of LFP activity in two frequency bands (1–5 Hz, 6–10 Hz), along with their corresponding reconstructions without shuffling, are shown in Figure 4C.

If reconstruction fidelity reflects sampling of the underlying population, it should scale with the number of neurons recorded. Consistent with this expectation, reconstruction accuracy increased with the number of neurons recorded per session for reconstructions preserving motif and opponent identity (Pearson r = 0.74, p=0.015; Figure 4D). No significant relationship between cell count and reconstruction accuracy was observed in motif-shuffled (r = 0.35, p = 0.32) or opponent-shuffled reconstructions (r = −0.36, p = 0.31; Supplementary Figure 5A). Finally, we asked whether reconstruction accuracy depended on the relative dominance of opponent motifs within each session. Reconstruction based on mean 1P calcium activity scaled strongly with the dominance of A motifs (Pearson r = 0.86, p = 0.0014; Figure 4E), with a similar dependence observed for motif-shuffled reconstructions (r = 0.67, p = 0.03; Supplementary Figure 5B). In contrast, reconstruction preserving motif and opponent identity was invariant to opponent balance (r = 0.11, p = 0.75).

### Spectral motifs are expressed in multi-frequency LFP dynamics

Given that LFP motif projections reliably tracked opponent ensemble activity, we asked whether the same spectral motifs could be recovered using LFP alone, without reference to calcium signals. To test this, we identified opponent motifs using LFP data alone by detecting patterns of amplitude co-fluctuations between all frequency bands (see Methods). This approach recovered the same set of opponent spectral motifs as those identified using LFP-calcium concordance (Supplementary Figure 6), indicating that motif structure is recoverable from multi-frequency LFP dynamics alone rather than being dependent on calcium measurements.

We then examined whether these spectral motifs were specific to mPFC or generalized across cortical regions. We applied the LFP-only motif detection procedure to retrosplenial cortex and anterior insula in a separate group of 11 rats (2–3 sessions each). Implant and recording procedures were adapted from a previous study^18^. The analyses revealed region-dependent structure: motif organization in anterior insula closely resembled that observed in mPFC, whereas retrosplenial cortex exhibited a distinct set of spectral motifs (Supplementary Figure 6). These results indicate that while opponent spectral motifs are not unique to a single cortical area, their specific spectral structure can vary across regions.

Finally, we asked whether opponent spectral motifs were tied to invasive recording modalities. Noninvasive recordings such as EEG provide an opportunity to test whether similar multiband structure can be detected noninvasively at larger spatial scales. Data were obtained from the Nencki-Symfonia EEG/ERP dataset^19,20^. Across 10 human subjects, we extracted spectral motifs from midline EEG electrodes (Fz, Cz, and Pz). Consistent with the rodent findings, opponent spectral motifs were recovered at each electrode site (Supplementary Figure 7). Together, these results indicate that opponent spectral motifs reflect a conserved organizational structure of multiband neural activity that generalizes across brain regions, species, and recording modalities, including noninvasive human EEG.

## Methods and materials

### Subjects

The study included 21 Long Evans rats (Charles River Laboratories). Six rats were used for fiber photometry experiments (3 male, 3 female), four rats were used for single-photon calcium imaging experiments (3 male, 1 female), and eleven rats were used for LFP-only analyses. Rats arrived at approximately one month of age (∼150 g) and were acclimated to the laboratory environment for ∼10-16 weeks prior to surgical implantation. Animals were housed in pairs in standard plastic tubs (10 × 10.75 × 19.5 in; Allentown, NJ, USA). Following surgery, rats were single-housed and received post-operative care consisting of analgesics (meloxicam for 3 days) and oral antibiotics (SMZ-TMP in drinking water for 5 days). Experiments were conducted approximately 4 weeks after surgery to allow for viral expression. All rats were maintained on a 12-h light/dark cycle with lights on at 6:00 A.M. Experiments were conducted during the light phase. Prior to recordings, rats were given *ad libitum* access to food and water. In the days preceding BCI training, rats were provided with 2 hours of unrestricted water access per day.

### Implant fabrication

#### Optrode

Fiber optic cannulae (1.25 mm outer diameter, 400 µm core diameter, 0.5 NA, 10 mm length; black ferrule, RWD) were secured to three 50 µm-diameter tungsten microwires (California Fine Wire) using black epoxy (J-B Weld). The tungsten wires were arranged along the dorsoventral axis with 400 µm spacing, with the most superficial wire positioned immediately above the fiber tip. All analyses were performed using signals from the first (most proximal) electrode.

#### Electrolens

A gradient-index (GRIN) lens (10 mm length) was secured with epoxy to five tungsten microwires using the same configuration as the optrode assembly. The microwires were arranged along the dorsoventral axis with 100 µm spacing between adjacent wires.

### Surgical Procedures

Stereotaxic surgeries were performed under aseptic conditions. Rats were anesthetized in an induction chamber with 5% isoflurane in room air using a low-flow anesthetic system (Kent Scientific) and maintained at 1.5-2.5% isoflurane via nose cone throughout the procedure. Body temperature was maintained at approximately 37 °C using a warming pad. Animals received a single preoperative dose of atropine (0.05 mg/kg), dexamethasone (0.5 mg/kg), and 0.9% saline (1 mL/kg, subcutaneous). Prior to incision, the scalp was shaved and sterilized with three alternating swabs of iodine and 70% ethanol. Bupivacaine (<0.2 mL, subcutaneous) was administered approximately one minute before incision.

After exposure and cleaning of the skull, burr holes were drilled using a 0.9 mm drill bit (Stoelting) to permit implant placement and secure anchoring skull screws (3-5 screws) as well as a ground/reference wire positioned several millimeters posterior to lambda over the cerebellum. For electrolens surgeries, a 3 mm diameter craniotomy was performed and centered over the implant coordinates.

For optrode experiments, 500 nL of AAV5-CaMKIIα-GCaMP6f-WPRE-SV40 (1.9 × 10¹³ GC/mL) was injected into mPFC (3.2 mm AP, 0.7 mm ML, 3.9 mm DV from skull, right hemisphere) using a 33-gauge Hamilton syringe mounted on a WPI microinfusion pump at a rate of 100 nL/min. For single-photon calcium imaging experiments, AAV1-CaMKIIα-GCaMP6m-WPRE-SV40 (3.8 × 10¹² GC/mL) was injected using the same infusion protocol (3.72 mm AP, 0.7 mm ML, 2.8 mm DV from brain surface, right hemisphere). Following completion of viral delivery, the syringe was left in place for 10 minutes before being slowly withdrawn.

Optrodes were implanted at 3.2 mm AP, 0.7 mm ML, and 3.5 mm DV from bregma in the right hemisphere and secured to the skull using C&B Metabond (Parkell Inc.) followed by black dental cement (AM Systems powder, Lang Dental resin). Tungsten wire leads were connected to a 32-channel EIB board (Neuralynx) using gold pins and further secured with dental cement. For electrolens implants (3.2 mm AP, 0.7 mm ML, 2.4 mm DV from brain surface, right hemisphere), Kwik-Sil (WPI) was applied to fill the gap between the lens and the brain prior to headcap construction using the same materials. One of the four animals implanted for single-photon imaging used a TDT ZCA-EIB32 ZIF-Clip adapter in place of the Neuralynx EIB board.

All surgical and experimental procedures adhered to NIH guidelines for the care and use of laboratory animals and were approved by the Institutional Animal Care and Use Committee at the San Diego VA Medical Center (Protocol A12-021).

### LFP processing

The raw LFP signal was acquired at 1017 Hz (RZ2, Tucker-Davis Technologies). For analyses, the signal was first notch filtered to remove line noise using a 60 Hz bandstop filter spanning 59–61 Hz. The signal was then bandpass filtered into overlapping 5 Hz frequency bins spanning 1–80 Hz (1–5 Hz, 2–6 Hz, …, 76–80 Hz) using zero-phase, 6th-order Butterworth filters to avoid phase distortion. For each band-limited signal, the analytic amplitude was extracted via a Hilbert transform. The resulting amplitude envelopes were then temporally smoothed with a 5-second Gaussian kernel with a standard deviation of 2.5 seconds and subsequently z-scored within session for normalization prior to further analyses.

### Fiber photometry processing

Fiber photometry signals were acquired at 1017 Hz with an online 20 Hz 6^th^ order low-pass filter applied during data collection. LED power was between 30-90 µW. Offline preprocessing was then performed on the 465 nm (calcium-dependent) and 405 nm (isosbestic control) channels, which were each first low-pass filtered at 5 Hz using zero-phase, 6th-order Butterworth filters. Photobleaching was corrected by fitting and subtracting a double-exponential function from each channel independently. To remove motion- and hemodynamic-related artifacts, the 405 nm signal was linearly fit to the 465 nm signal and subtracted, yielding a corrected calcium-dependent fluorescence trace. The resulting signal was subsequently low-pass filtered at 0.1 Hz using a zero-phase, 6th-order Butterworth filter to isolate slow population-level calcium dynamics and z-scored within session prior to analysis.

### Calcium Imaging Preprocessing and CNMF-E Extraction

Miniscope videos were recorded at 20 Hz (Inscopix nVoke), and subsequently spatially and temporally downsampled by a factor of three. Frames exhibiting excessive motion were manually excluded. Images were then spatially filtered to suppress low-frequency background structure, followed by motion correction across remaining frames. Neuronal sources were then identified using constrained nonnegative matrix factorization for endoscopic data (CNMF-E) with the following parameters: expected cell diameter of 12 pixels, minimum spatial correlation of 0.92, merge threshold of 0.92, patch size of 100 pixels with 30-pixel overlap, ring size factor of 2 for local background estimation, and a minimum signal-to-noise ratio (SNR) of 10. Extracted components were manually curated prior to downstream analysis.

### CNMF-E Trace Processing

CNMF-E–extracted calcium traces were loaded from CSV files and restricted to accepted components based on manual acceptance flags. CNMF-E traces and inferred spike trains were temporally aligned using shared GPIO synchronization pulses and truncated to matched recording epochs. Cells with no detected spiking activity were excluded from analysis.

All calcium traces were linearly interpolated to a common time base for alignment with electrophysiology recordings. Global fluorescence fluctuations were estimated from whole-field averages extracted from raw videos to which motion correction parameters learned from the spatially filtered data had been applied. The average fluorescence was detrended using a double-exponential fit, low-pass filtered at 1 Hz using zero-phase filtering, and z-scored. Cell-wise calcium signals were also low-pass filtered at 1 Hz to isolate slow population-level calcium dynamics.

Calcium spike trains were inferred via deconvolution (fudge factor = 0.99; spike SNR threshold = 5). Deconvolution event timestamps were mapped to imaging frames, and events were additionally filtered by retaining only those whose coincident CNMF-E fluorescence exceeded the cell’s mean CNMF-E level. For analyses requiring a rate-like signal, binary event trains were converted to a smoothed event-rate estimate by applying a 10 s moving-average window and scaling by the imaging sampling rate (yielding units of events/s). All primary results were invariant to variations in spike-detection thresholds and smoothing window length.

### ROI Extraction

Binary spatial masks for each accepted CNMF-E component were loaded from cell-specific image files and thresholded to produce binarized cell footprints. These pixel index sets were stored as individual region-of-interest (ROI) masks. ROIs were represented as binary images in which pixels belonging to the accepted cell footprint were assigned a value of 1 and all other pixels were set to 0. Only ROIs corresponding to manually accepted components were retained for downstream analyses.

### Spectral Motif Identification

For each time point, a signed coupling vector was constructed across 76 overlapping 5 Hz frequency bands spanning 1-80 Hz by taking the sign of the product between each band’s z-scored smoothed amplitude envelope and the z-scored calcium activity. This yielded a discrete, multiband representation indicating whether activity in each frequency band was positively or negatively aligned with the calcium activity at each sample. Similarity between time points was quantified using correlation distance between their signed multiband vectors, and average-linkage hierarchical clustering was applied to group time points with similar frequency-alignment structure. Clusters were then iteratively consolidated into candidate motifs using a projection-based similarity criterion. For each cluster, a provisional motif was defined as the mean signed pattern across its member time points. Time points were projected onto these provisional motifs, and clusters were merged only if the correlation between their projection vectors exceeded a predefined threshold (0.4-0.8). This consolidation procedure was repeated until the number and composition of clusters stabilized, followed by reclustering on the resulting motifs to ensure convergence. This process was iterated for each threshold.

To assess motif robustness and control for autocorrelation-driven structure, a null distribution was generated by independently circularly shifting each frequency band by random offsets drawn from multiples of 20 seconds. The full clustering and consolidation procedure was repeated on each shifted dataset, yielding an empirical distribution of maximum cluster sizes expected by chance. In the unshifted data, only motifs whose membership exceeded this null distribution were retained. Motifs were additionally required to be expressed across all sessions and animals, ensuring that identified patterns reflected consistent, cross-recording structure rather than session-specific effects.

To reduce sensitivity to any single consolidation threshold, we repeated motif identification across thresholds (0.4-0.8). Within each threshold, candidate motifs were filtered using a shift-based null distribution and required to be expressed across all sessions and animals. We then pooled motif assignments across thresholds and computed final motif templates by averaging signed coupling vectors across all time points assigned to each motif across threshold runs, yielding threshold-robust (hyperparameter-agnostic) estimates. Motifs were organized into canonical opponent families by PCA on motif templates, assigning each motif to its dominant component and splitting by loading sign; each class was then refined by excluding time points assigned to other classes and recomputing the mean pattern.

### Motif Projections

To quantify motif expression, two complementary projection measures were computed. Concordance-based motif projections were defined by first binarizing each frequency band and the calcium activity by sign (positive vs negative deviation), yielding a signed agreement vector that encoded whether each LFP band was aligned or anti-aligned with the calcium signal at each time point. These signed concordance vectors were then projected onto the motif templates to measure the degree to which instantaneous cross-modal alignment matched each motif. In contrast, LFP projections were computed using binarized LFP activity alone, independent of calcium, by binarizing each frequency band by calculating the sign of its z-scored smoothed amplitude (above vs below baseline) and projecting these bandwise binary patterns onto the same motif templates. Thus, while both projections quantify motif expression in the same motif space, the concordance-based projection captures coordinated sign agreement between LFP and calcium, whereas the LFP projection reflects multiband LFP structure without reference to cross-modal coupling.

### Brain-Computer Interface Training

BCI training was implemented as a closed-loop BCI task controlled in real time via Synapse and RPvdsEx (RZ2 system, Tucker-Davis Technologies) and a MATLAB controller that interfaced with these programs via UDP communication. Online signal evaluation was performed in fixed 100 ms bins. Task performance was mapped to an auditory feedback tone whose frequency was updated every bin as a monotonic function of the control signal relative to a threshold. In sessions where animals were trained to decrease the control signal, tone frequency ranged from 6-12 kHz, with trials starting at 6 kHz and increasing in frequency as each bin that was lower than the threshold. Bins above the threshold decreased the tone pitch towards 6 kHz, which served as the minimum possible frequency. Changes in tone frequency were incremented such that perfect performance would move the cursor from 6 kHz to 12 kHz in 2 seconds. Tone presentation duration was fixed at 2 s at trial onset for BCI trials to initiate each trial. Trials terminated when the frequency of the tone reached 12 kHz (successful trials), or after 10 seconds elapsed (unsuccessful trials). Training was conducted over approximately one month and consisted of interleaved BCI trials and control trials. During BCI trials, tone frequency provided continuous performance-contingent feedback. Successful trials triggered delivery of a water reward for a fixed duration (1.5 s), followed by a fixed inter-trial interval of 15 s. In contrast, control trials provided no auditory feedback or reward.

Feedback thresholds were initialized either from the most recent prior session or via an online calibration procedure when no prior threshold was available. Calibration was implemented as an adaptive binary search, in which control-trial outcomes were accumulated in blocks of 10 trials, and the threshold was adjusted upward or downward depending on whether performance exceeded or fell below a target success rate of 60%. During BCI training, threshold adaptation was driven exclusively by control trials to stabilize task difficulty independently of learning-related changes during BCI trials. Threshold updates during training were applied every 5 completed control trials, scaled by the deviation of recent control-trial performance from the target success rate and by the direction of training (increase vs. decrease of the control signal). Sessions terminated upon reaching a predefined maximum duration or trial count, and all task parameters, outcomes, and threshold trajectories were stored for offline analysis.

### Histology

To confirm viral expression and implant location, rats were perfused with 0.9 % saline followed by 4 % paraformaldehyde (PFA). Upon extraction, the brains were held in 30% sucrose solution for at least 2 days and cryoprotected. Each brain was sectioned at 50 μm thickness using a cryostat (Leica CM1950). Sections were counterstained with DAPI and analyzed under a confocal microscope (Zeiss LSM 880).

### Statistical analyses

All analyses were performed in MATLAB. Unless otherwise stated, statistics were computed at the session level, with metrics first calculated within session and then aggregated across sessions and animals. Normality was not assumed a priori. Parametric or nonparametric tests were selected based on comparison structure. All tests were two-sided, and exact p-values and sample sizes are reported in the text and figures.

Relationships between LFP activity, calcium signals, motif expression, and behavior were quantified using Pearson correlations. When comparing correlation strength across conditions or models, coefficients were Fisher z-transformed prior to statistical testing. Paired comparisons within sessions (e.g., motif A vs. motif B dominance, early vs. late learning, BCI vs. control trials) were evaluated using paired t-tests or Wilcoxon signed-rank tests, as appropriate. Comparisons between groups with unequal variance were assessed using Welch’s t-tests. One-sample t-tests were used to test whether session-level summary metrics differed from zero.

For analyses involving multiple frequency bands or motifs, significance was assessed using empirically derived null distributions rather than parametric multiple-comparison corrections. Motif robustness was evaluated using permutation tests in which individual frequency bands were circularly shifted by random temporal offsets, preserving marginal statistics and autocorrelation structure. Observed motif sizes and projection strengths were compared against these null distributions, and only motifs exceeding null expectations were retained.

Spatial organization of motif-defined neuronal ensembles was assessed using nearest-neighbor analyses of ROI centroids. For each neuron, the identity of its closest spatial neighbor was determined, and motif-to-motif neighbor fractions were computed. Cells belonging to motifs with fewer than five neurons per session were excluded. Null distributions were generated using 10,000 permutations per session in which motif labels were randomly reassigned while preserving cell counts and spatial locations. Empirical p-values were computed as the proportion of shuffled values greater than or equal to the observed value, with a +1 correction applied to avoid zero p-values.

Reconstruction accuracy and decoding performance were compared across conditions using paired t-tests on Fisher z–transformed correlations. Control reconstructions were generated by shuffling motif identity or opponent assignment while preserving marginal statistics. Relationships between reconstruction accuracy, cell count, and motif dominance were assessed using Pearson correlation across sessions.

All summary values are reported as mean ± SEM unless otherwise stated. Sample sizes are provided in the figure legends. No statistical methods were used to predetermine sample size.

## Acknowledgements

Burroughs Wellcome Fund, 1015644

National Institute of Mental Health, R01MH123650

Hope for Depression Research Foundation, 30063773

---

**Supplementary Figure 1.**
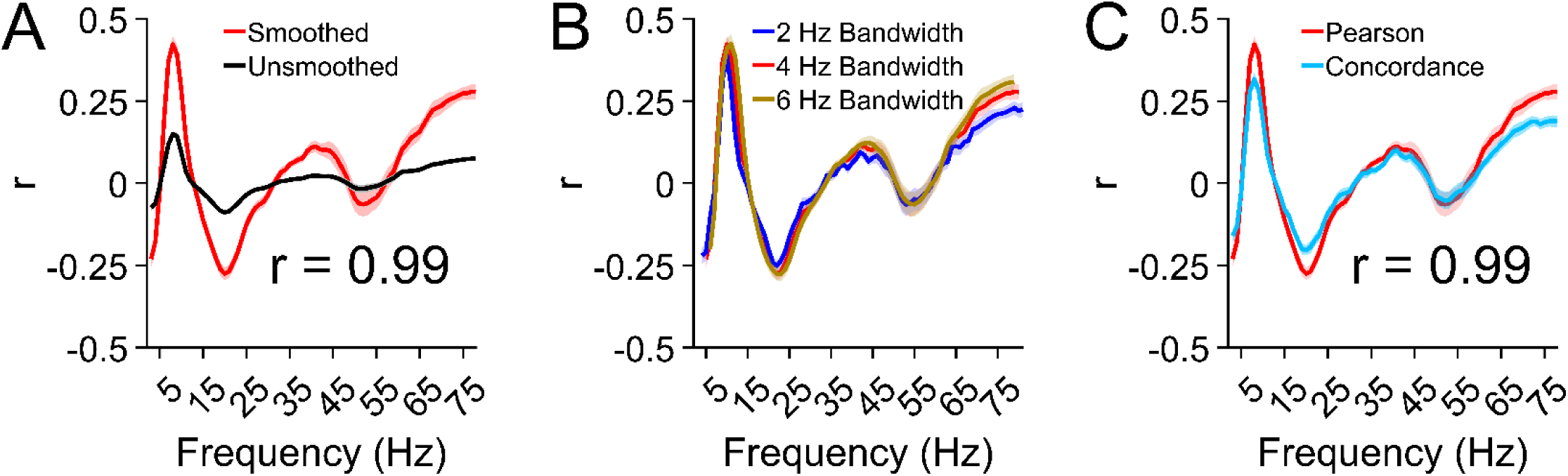
Frequency–calcium relationships are robust to amplitude smoothing and bandwidth definition. (A) Comparison of smoothed and unsmoothed band-limited LFP amplitude envelopes used to compute frequency-calcium relationships. (B) Frequency-specific correlations computed using different bandwidth definitions (2 Hz, blue; 4 Hz, red; 6 Hz, gold), demonstrating preservation of spectral structure across bandwidth choices. (C) Frequency-calcium relationships quantified using Pearson correlation (red) and mean concordance (light blue, see Methods), showing consistent spectral organization across metrics.

**Supplementary Figure 2.**
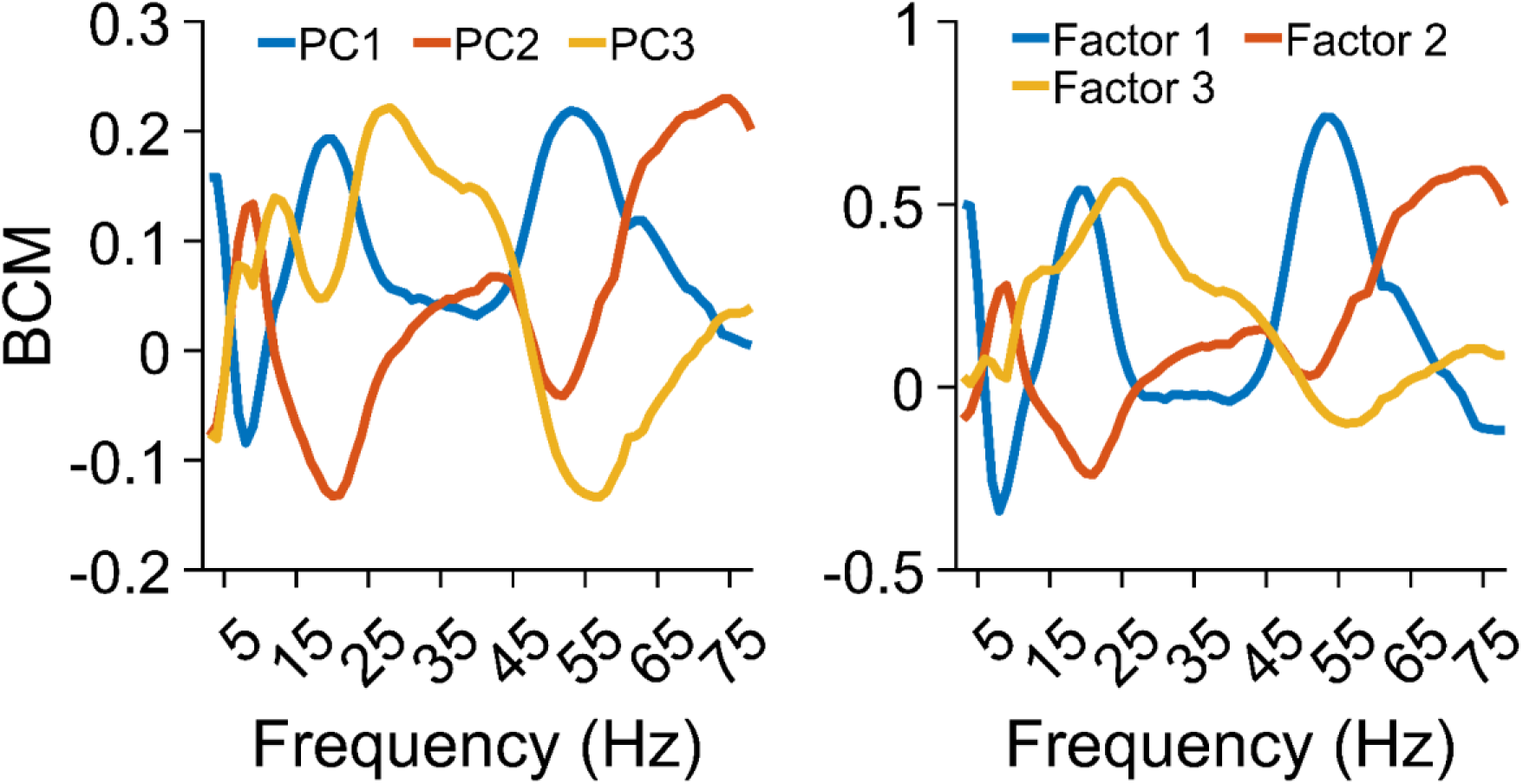
Opponent spectral motif structure is preserved across factorization methods. Recurring multiband amplitude configurations identified directly from band-limited LFP concordance with calcium activity (see Methods) using PCA and factor analysis.

**Supplementary Figure 3.**
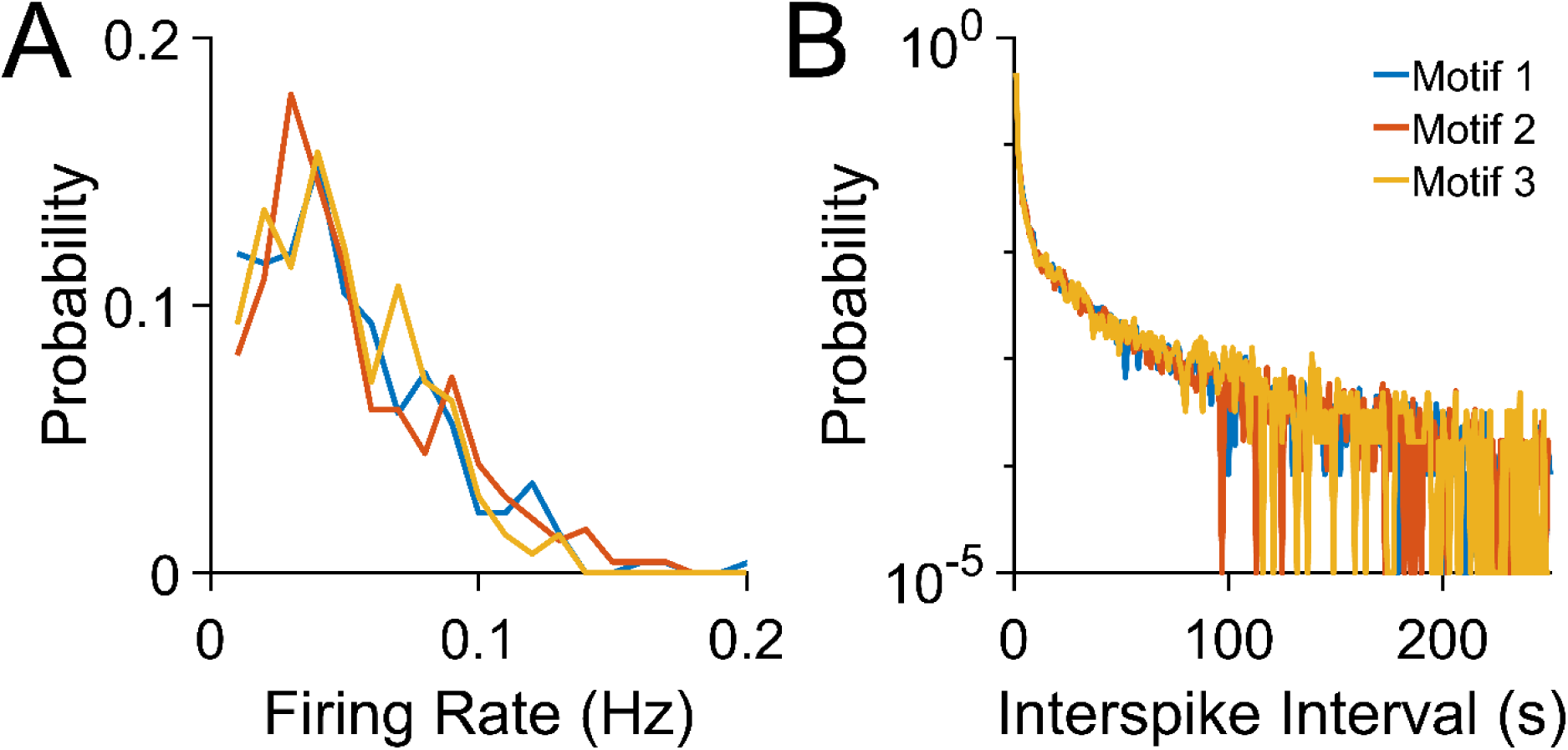
Motif-defined ensembles are not distinguished by gross firing statistics. (A) Distribution of mean firing rates for neurons assigned to each motif. No consistent differences were observed across sessions. (B) Interspike interval (ISI) distributions for neurons grouped by opponent motif assignment. ISI structure did not systematically differ across motifs.

**Supplementary Figure 4.**
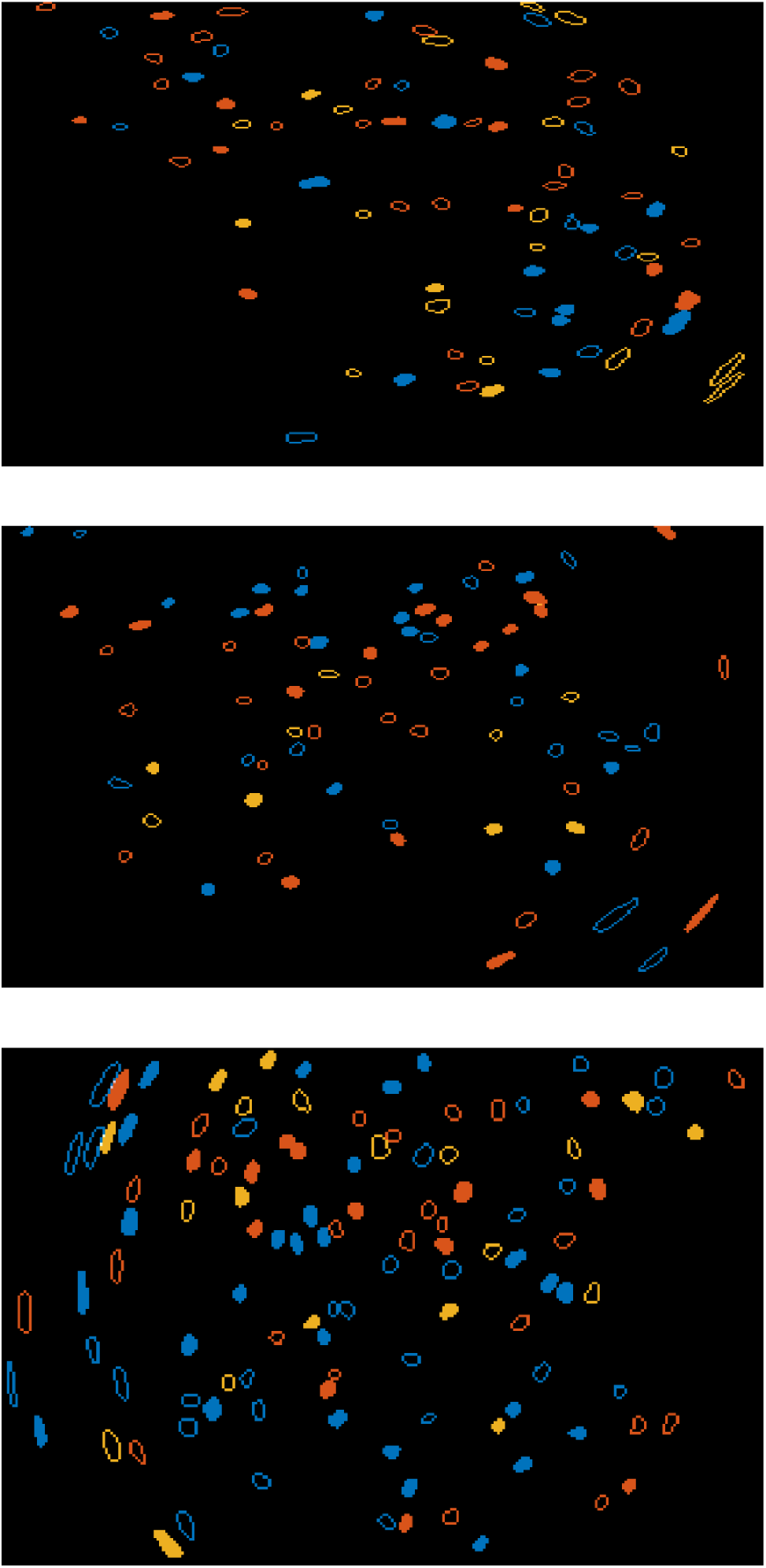
Opponent neuronal ensembles are spatially intermingled within mPFC. Spatial distribution of neurons assigned to Motif 1 (blue), Motif 2 (orange), and Motif 3 (yellow) across three sessions from three rats. Open and closed markers denote opposing A and B components within each motif. Neurons belonging to opponent components were interspersed throughout the imaging field rather than anatomically clustered. Nearest-neighbor permutation analyses revealed minimal deviation from chance organization, with small effect sizes and few significant motif-pair comparisons relative to shuffled controls (97.6% of permutations p > 0.05), indicating that motif-defined ensembles reflect functional rather than anatomical segregation.

**Supplementary Figure 5.**
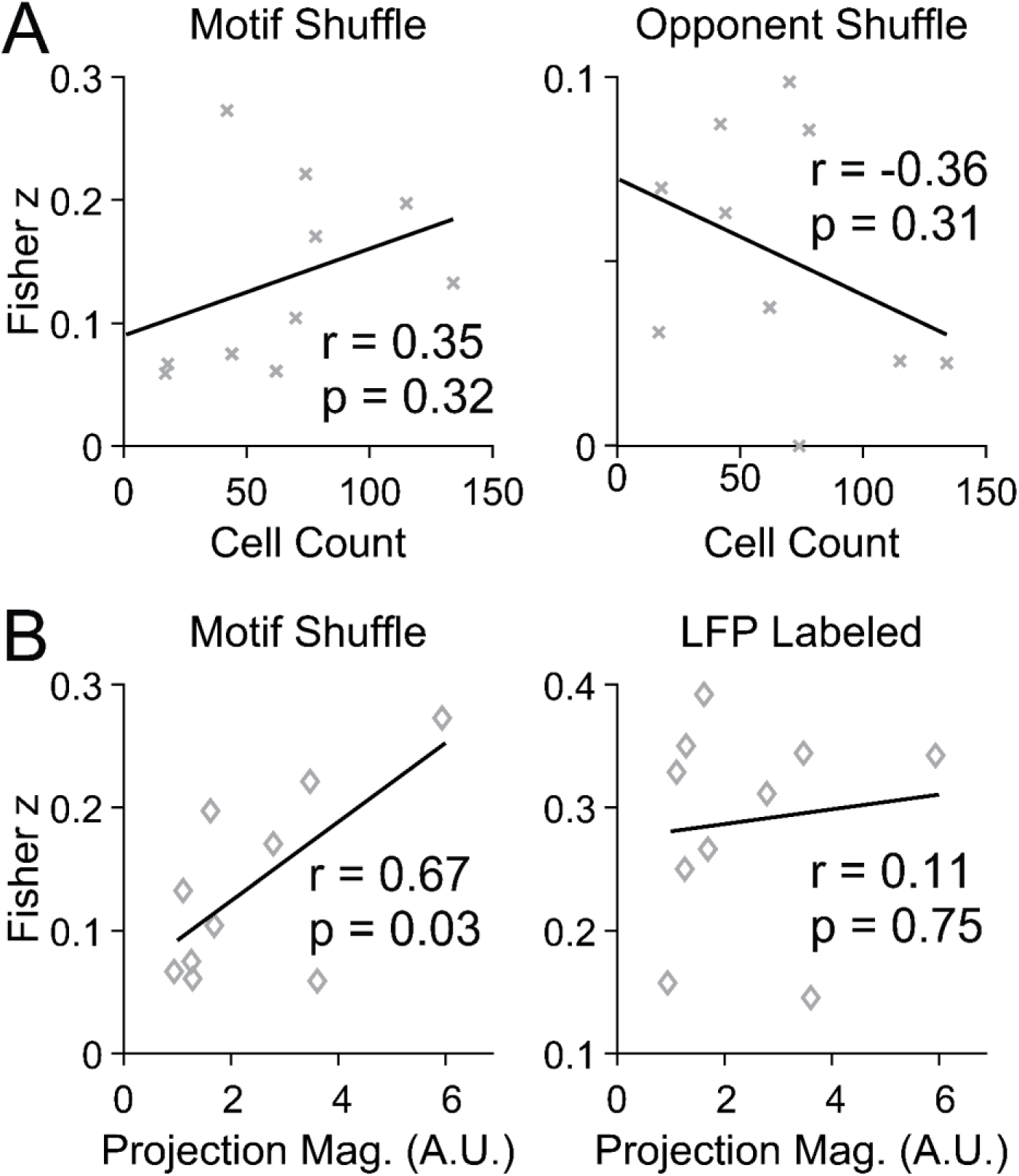
Control reconstructions fail to scale with cell count and remain dependent on opponent motif dominance. (A) Relationship between reconstruction accuracy and number of recorded neurons per session for control models. Motif-shuffled (left) and opponent-shuffled (right) reconstructions did not scale with cell count (motif-shuffled: r = 0.35, p = 0.32; opponent-shuffled: r = −0.36, p = 0.31). (B) Dependence of reconstruction accuracy on relative dominance of A versus B opponent motifs. Motif-shuffled reconstructions showed a dependence of A-motif dominance (r = 0.67, p = 0.03, left). In contrast, reconstructions preserving motif and opponent identity were invariant to opponent balance (r = 0.11, p = 0.75, right), indicating that accurate reconstruction depends on opponent structure rather than global shifts in ensemble activity.

**Supplementary Figure 6.**
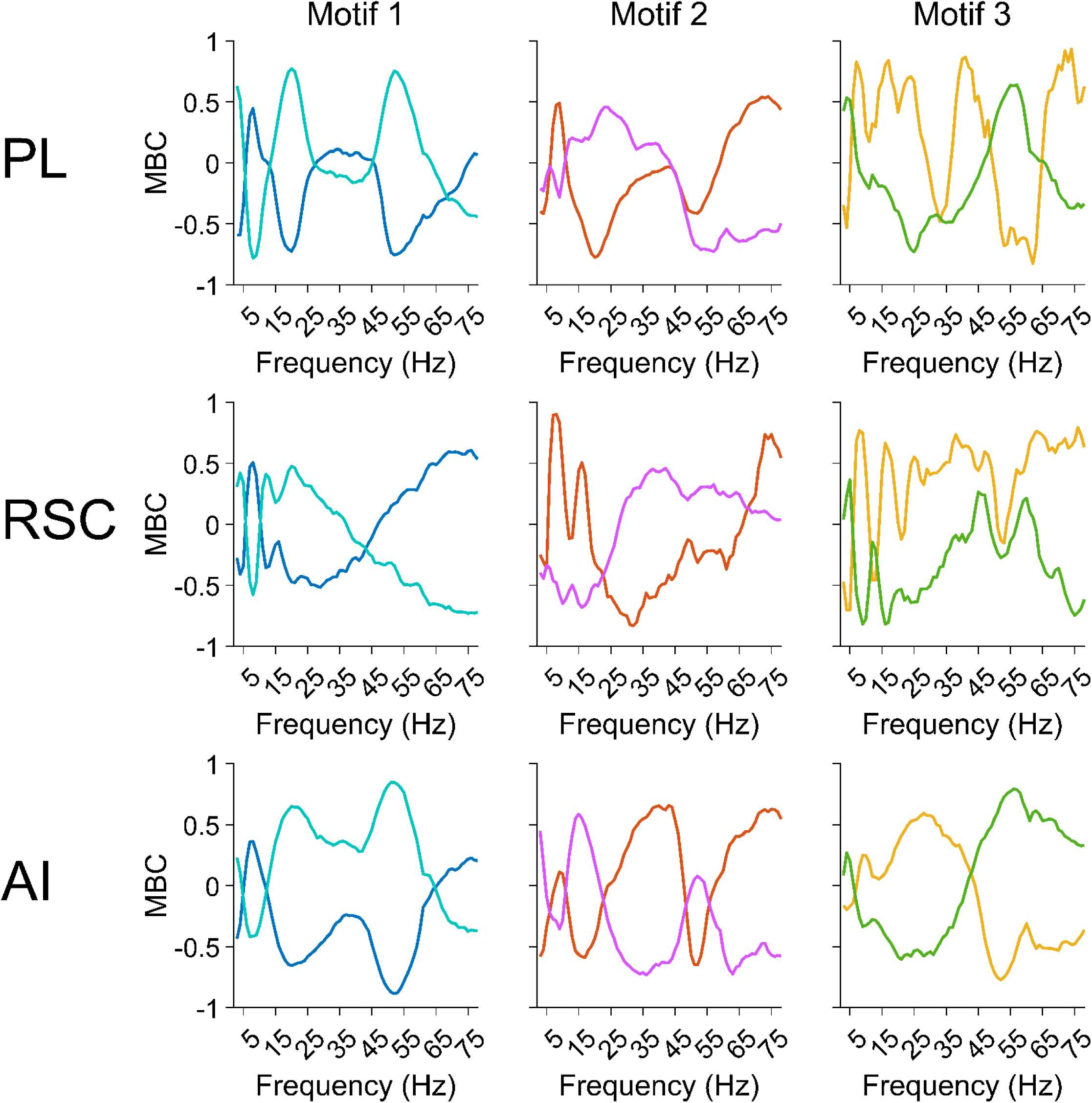
Opponent spectral motifs are recoverable from multiband LFP dynamics and vary across cortical regions. (Top) Opponent spectral motifs identified using LFP data alone by clustering multiband amplitude co-fluctuations without reference to calcium activity. (Middle) LFP-only motif extraction in RSC. (Bottom) LFP-only motif extraction in AI.

**Supplementary Figure 7.**
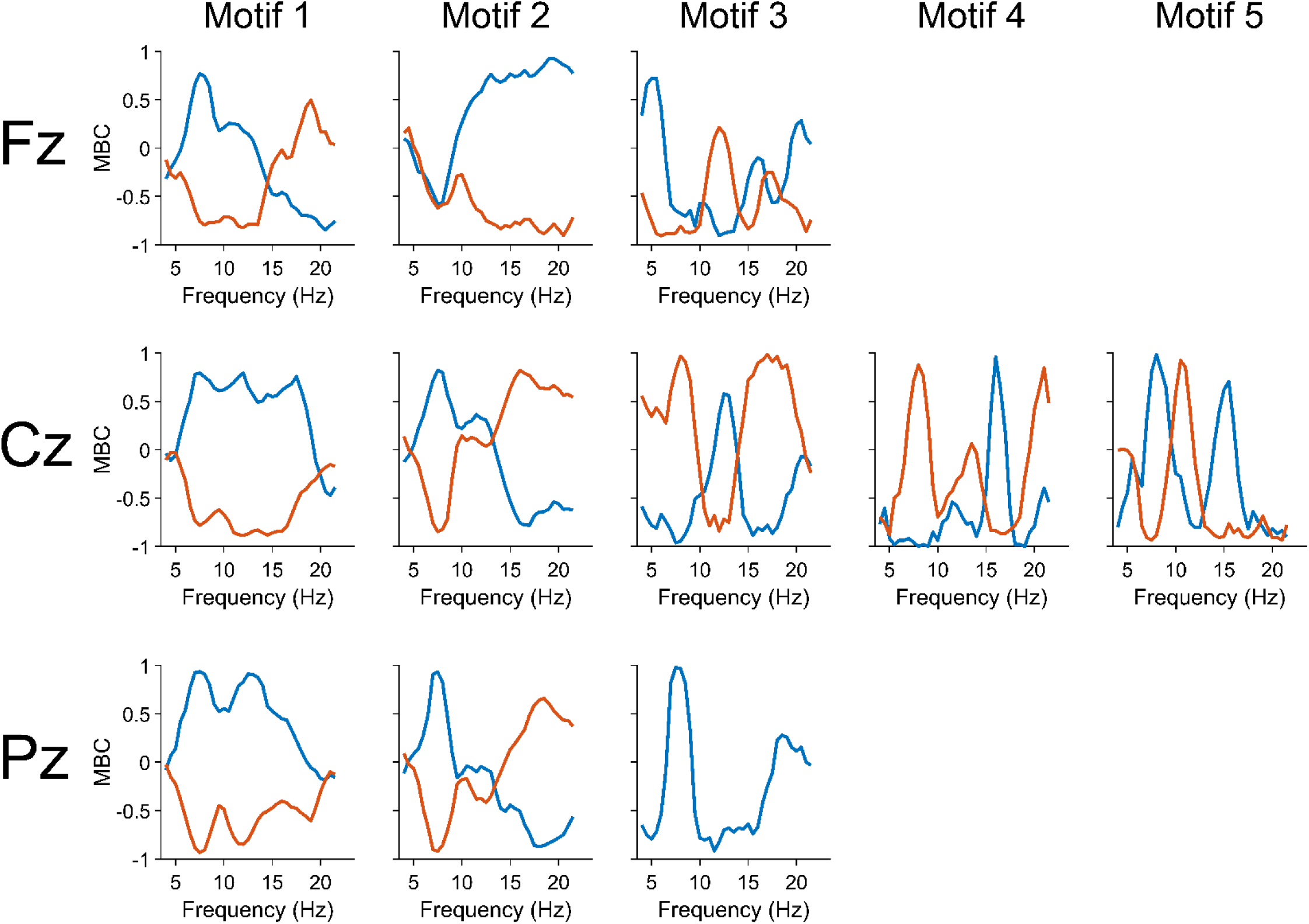
Opponent multiband spectral motifs generalize to noninvasive human EEG. Spectral motifs were extracted from midline EEG electrodes (Fz, Cz, and Pz) across 10 human subjects using the same multiband amplitude co-fluctuation procedure applied to rodent LFP recordings. Motif structure was reproducible across subjects and electrode sites, and recapitulated opponent dynamics observed in rats.

